# diaPASEF analysis for HLA-I peptides enables quantification of common cancer neoantigens

**DOI:** 10.1101/2024.07.30.605861

**Authors:** Denys Oliinyk, Hem Gurung, Zhenru Zhou, Kristin Leskoske, Christopher M. Rose, Susan Klaeger

## Abstract

Human leukocyte antigen class I (HLA-I) molecules present short peptide sequences from endogenous or foreign proteins to cytotoxic T cells. The low abundance of HLA-I peptides poses significant technical challenges for their identification and accurate quantification. While mass spectrometry (MS) is currently a method of choice for direct system-wide identification of cellular immunopeptidome, there is still a need for enhanced sensitivity in detecting and quantifying tumor specific epitopes. As gas phase separation in data-dependent MS data acquisition (DDA) increased HLA-I peptide detection by up to 50%, here, we aimed to evaluate the performance of data-independent acquisition (DIA) in combination with ion mobility (diaPASEF) for high-sensitivity identification of HLA presented peptides. Our streamlined diaPASEF workflow enabled identification of 11,412 unique peptides from 12.5 million A375 cells and 3,426 8-11mers from as low as 500,000 cells with high reproducibility. By taking advantage of HLA binder-specific *in-silico* predicted spectral libraries, we were able to further increase the number of identified HLA-I peptides. We applied SILAC-DIA to a mixture of labeled HLA-I peptides, calculated heavy-to-light ratios for 7,742 peptides across 5 conditions and demonstrated that diaPASEF achieves high quantitative accuracy up to 4-fold dilution. Finally, we identified and quantified shared neoantigens in a monoallelic C1R cell line model. By spiking in heavy synthetic peptides, we verified the identification of the peptide sequences and calculated relative abundances for 13 neoantigens. Taken together, diaPASEF analysis workflows for HLA-I peptides can increase the peptidome coverage for lower sample amounts. The sensitivity and quantitative precision provided by DIA can enable the detection and quantification of less abundant peptide species such as neoantigens across samples from the same background.

## Introduction

Sensitive mass spectrometry (MS) methods have greatly advanced the discovery of peptides presented by major histocompatibility complex (MHC) or human leukocyte antigen (HLA) molecules and helped inform cancer immunotherapies(Yewdell 2022,Bassani-Sternberg 2016,Yadav 2014). However, low abundance, poor recovery yields, and lack of clear digestion rules pose significant challenges for efficient identification of HLA-I peptidome.

As of today, data-dependent acquisition (DDA) remains a method of choice for MS-based immunopeptidomics. High quality of acquired MS/MS spectra obtained in DDA mode can be readily translated into high confidence determination of peptide sequences that makes DDA ideal for discovery immunopeptidomics. Nevertheless, DDA inherently suffers from stochastic intensity-based selection of precursor ions that often leads to low sensitivity and low data completeness. Data-independent acquisition (DIA), in contrast to DDA, employs fragmentation of all precursor ions that fall into predefined mass windows, generating highly complex fragment spectra. Fragmentation of all available precursor ions without prior selection significantly enhances sensitivity and data completeness of an analysis. In recent years, DIA has proven to be an attractive acquisition method for label free quantification in global proteomics. Analysis of highly complex DIA fragment spectra is a non-trivial task, which traditionally employs time and resource intensive DDA-based spectral libraries. Development and rapid evolution of advanced computational tools enabled an accurate prediction of peptide retention time, ion mobility, and fragment ion intensity(1–5). This information can then be parsed into *in-silico* predicted spectral libraries that can serve as an attractive alternative to empirical spectral libraries.

While DIA for immunopeptidomics is not yet widely adapted, more groups have been exploring this type of acquisition for HLA peptides. For example, DIA has been used for identification of neoantigens (6) or for identification of peptides bound to soluble HLA (sHLA)(7). A significant drawback for use of DIA in immunopeptidomic applications stems from the lack of clear digestion rules for HLA-presented peptides which makes the generation of predicted spectral library a computationally challenging task. Therefore, immunopeptidomics mainly relies on generation of experimental spectral libraries. This is disadvantageous if analyzing samples across multiple cell lines or patients with varying HLA types as each of these will present peptides with different binding rules and varying amino acid anchor residues. Moreover, the nonspecific nature of these peptides also complicates data analysis as co-eluting precursors might lead to false identifications.

We have recently shown that ion mobility separation, provided by high field asymmetric waveform ion mobility (FAIMS, Thermo Fisher)(8) or trapped ion mobility spectrometry (tims)(9), helped to increase HLA-I and HLA-II peptide identification starting from more reasonable sample input amounts. Increased sensitivity due to gas phase separation provided by these approaches facilitate detection of substoichiometric post translational modifications (10) or neoantigens.

Here, we evaluate the quantitative accuracy of DIA for HLA Class I peptides in combination with ion mobility on the timsTOFscp and propose an *in silico* library approach using a combination of prediction tools. We then used DIA for identification and quantification of previously identified shared neoantigens in an engineered monoallelic cell line model.

## Materials and Methods

### Cell culture and SILAC labeling

A375 cells were cultured in DMEM supplemented with 10% FBS, 2 mM L-glutamine and 1% Penicillin-Streptomycin (Gibco). For SILAC experiments, A375 cells were passaged at least 7 times in powdered DMEM for SILAC (Thermo Scientific) prepared with either light or heavy amino acid isotopes (84 mg/L L-arginine:HCl [^13^C_6_,^15^N_4_], 175 mg/L L-lysine:2HCl [^13^C_6_,^15^N_2_], 100 mg/L L-leucine [^13^C_6_]; Cambridge Isotope Laboratories, Inc.), 44 mM sodium bicarbonate, 10% dialyzed FBS (Thermo Scientific), 200 mg/L L-proline, 2 mM L-glutamine, and 1% Penicillin-Streptomycin (Gibco). Cells were grown to 90% confluency, harvested with Trypsin-EDTA, washed three times with ice cold PBS and snap frozen.

C1R HLA-A*11:01 monoallelic line containing 47 common cancer mutations was engineered as previously described (Gurung 2023). Briefly, HLA-I null C1R cells were electroporated with a piggyBac neoantigen expression plasmid system and transduced with lentiviral HLA expression vector. Cells were grown in IMDM supplemented with 10% FBS and 1ug/ml puromycin (Gibco).

### HLA-I Peptide Enrichment and Peptide Elution

A375 cell pellets were lysed in lysis buffer containing 1% CHAPS, 20mM Tris, pH 8.0, 15mM NaCl, 2mM MgCl2, 0.2 mM iodoacetamide, 1mM EDTA, 1x Complete Protease Inhibitor Tablet-EDTA free, 2µl benzonase, 0.2 mM PMSF in total of 2 ml lysis buffer per 100 million cells. Each lysate was incubated on ice for 30 min and vortexed every 5 min. Lysates were then centrifuged at 20,000 rcf for 15 min at 4°C and supernatants were transferred to 1 ml 96 deep-well plates. For neoantigen detection, a 500 million cell pellet of a C1R A*11:01 monoallelic cell line containing stably transfected 47-mer cassette without GS linkers was lysed in 5 mL 1% CHAPS lysis buffer and appropriate volume was taken for the respective dilution point for HLA Class I enrichment.

Immunoprecipitation (IP) of HLA-I peptides and sample preclear was performed on AssayMAP Bravo Sample Prep Platform (Agilent), using Affinity Purification v.4 Protocol. Sample preclear was performed by dispensing lysates through 5 µl Protein-A cartridges (Agilent), previously primed and equilibrated with PBS. For enrichment of HLA-I complexes, flow through was loaded on 5 µl W6/32 cross linked Protein-A cartridges. Cartridges were primed and equilibrated with 100 µl of 20mM Tris pH 8.0 and 150mM NaCl in water. After sample loading, cartridges were washed with 250 µl of 20 mM Tris pH 8.0 and 400mM NaCl in water followed by final wash with 100 µl of 20mM Tris pH 8.0 in water. The antibody-bound HLA-peptide complexes were eluted with 60 µl of 0.1M acetic acid in 0.15% trifluoroacetic acid (TFA) and collected to 96 well PCR, Full Skirt, PolyPro plates (Eppendorf). 2 µl of 30% NH4OH was added to the eluate to neutralize the pH. Samples were then reduced with 5 mM Dithiothreitol (DTT) at 56°C for 15 min, and alkylated with 10 mM Iodoacetamide (IAA) in the dark at RT for 20 min. The samples were then acidified by adding 10% TFA to pH ~ 3.

Next, samples were loaded on the AssayMAP Bravo platform for C18-based desalting. 5 µl C18 cartridges (Agilent) were primed with 80% acetonitrile (ACN) 0.1% TFA and equilibrated with 0.1% TFA. The samples were afterwards loaded through the cartridges, washed with 0.1% TFA and eluted with 30% ACN in 0.1% TFA and dried by vacuum centrifugation.

### LC-MS/MS Analysis

Purified and desalted peptides were reconstituted in 4 µl of solvent A (0.1% formic acid) and separated via nanoflow reversed-phase liquid chromatography (Vanquish Neo, Thermo) with 60 min or 120 min methods at flow rate of 0.3 µl/min on an Aurora Ultimate nanoflow UHPLC column with CSI fitting (25 cm x 75 μm ID, 1.7 μm C18) (IonOpticks) at 50°C. Biognosys iRT peptides were spiked into each sample for both DDA and DIA acquisitions. For HLA-A*11:01 dilution DIA runs 100 fmol of previously targeted neoantigens (Gurung 2023) were spiked into the samples. Mobile phase A was water with 0.1 vol% formic acid and B was ACN with 0.1 vol% formic acid. Peptides eluting from the column were electrosprayed (CaptiveSpray) into a TIMS quadrupole time-of-flight mass spectrometer (Bruker timsTOF SCP). When operated in dda-PASEF, a ten or three PASEF/MSMS scan per topN acquisition method was utilized. An adapted polygon in the m/z and ion mobility space was employed(9, 11). The mass spectrometer was operated with an accumulation and ramp time of 166 ms or 300 ms. Precursors were isolated with a 2 Th window below m/z 700 and 3 Th above and actively excluded for 0.4 min with a threshold of 10,000 arbitrary units. A range from 100 to 2000 m/z and 0.65 to 1.67 Vs cm-2 was covered with a collision energy 20 eV at 0.6 Vs cm-2 with a linear increase to 59 eV at 1.6 Vs cm-2. When operated in dia-PASEF mode, an optimized isolation window scheme in the m/z vs ion mobility plane was designed using pyDIAid(12), covering >99.5% of all precursor ions including singly charged species. The method covered precursors within 300-1200 Da. The mass spectrometer was operated in sensitivity mode with an accumulation and ramp time of 100 ms and a cycle time of 1.17 s.

### Raw data analysis

DDA data was analyzed using FragPipe v20 with a nonspecific HLA workflow, peptide length was set to 7-15 mers against a human database (Uniprot 08/2023, 104,452 protein sequences) including swissprot and trembl entries and 48 common contaminants. Precursor and fragment mass tolerance were both set to 20 ppm. Carbamidomethylation of cysteines was set as static modification and methionine oxidation, cysteinylation, and pyro-Glu or acetylation of the N terminus were set as variable modification. We allowed a maximum of three variable modifications per peptide. For validation, MSBooster in combination with Percolator was enabled and protein FDR was disabled. A spectral library was built with the spectral library module in FragPipe using standard parameters and retention time alignment to iRT standards (Biognosys).

DIA data was analyzed using DIA-NN version 1.8.2 beta 11 with standard settings, no match between runs, precursor FDR filtering at 1% searching against either sample specific spectral library, generated by FragPipe, or a predicted library.

SILAC DIA data was analyzed with DIA-NN version 1.8.2 beta 11 with following search settings: Library Generation was set to “IDs, RT and IM Profiling”, Quantification Strategy was set to “Peak height”, Scan window = 1, Mass accuracy = 10 p.p.m. and MS1 accuracy = 5 p.p.m. Additional commands, entered in DIA-NN command window: {--peak-translation}, {--fixed-mod SILAC,0.0,LKR,label}, {--lib-fixed-mod SILAC}, {--channels SILAC,L,LKR,0:0:0; SILAC,H,LKR,6.020129:8.014199:10.008269}.

### HLA Peptide Binding Prediction

HLA peptide to allele binding was predicted with HLApollo v1. To evaluate MS peptide identifications, identified peptide sequences of length 8-13 were evaluated for binding on the cell lines’ respective alleles. Peptides were assigned as binders with an HLApollo score >0(13), peptides predicted to bind to multiple alleles were assigned to every possible allele. For spectral library generation, all possible 8-13mers from the entire human proteome (swissprot: 62,266 protein sequences, 57,172,493 possible 8-13mer peptides including sequence context for 4 residues at the N and C terminus) were generated and predicted for presentation on A375 cells. Peptides were only retained if they would bind to any of the six alleles with a score >0.

### Peptide Retention Time, Fragmentation and Ion Mobility Prediction with Salud and Ionmob

For the predicted A375 spectral library (predicted library), HLA peptides identified as binders by HLApollo were considered for charge states 1-3 per peptide with fixed cysteine alkylation and variable methionine oxidation as possible modifications. Spectral features were predicted using Salud, an in-house collection of ML models based on Prosit(4) to predict retention time and fragmentation. The peptide fragmentation model was based on the Prosit nontryptic HCD model(14) and the collision energies used for prediction were 15, 20, and 25 for charge states 1, 2, and 3, respectively. For ion mobility, collisional cross sections were predicted using Ionmob(5) and converted to 1/K_0_ values.

### Neoantigen identification and quantification

In order to systematically identify HLA A*11:01 neoantigens, a two step approach was used to build a spectral library (A1101-combo) in Fragpipe v20. First, a spectral library was created from all A*11:01 DDA runs using a human proteome fasta file containing the neoantigen sequences along with BFP reporter sequences and iRT sequences (Biognosys). In order to maximize chances of identifying the putative neoantigens, three replicate DDA injections of an equimolar mix of 456 unique synthetic heavy 8-11mer neoantigens that potentially resulted from 47 common cancer mutations (15) were performed as described above using an effective gradient of 51 minutes. Since the synthetic neoantigen pool contained one heavy amino acid per peptide, mass delta for lysine (K: 8.01499), arginine (R: 10.008269), valine/proline (V/R: 6.0138), leucine/isoleucine (L/I: 7.0171), phenylalanine/tryptophan (F/Y: 10.0272), and alanine (A:4.007) along with methionine oxidation were set as variable modifications. We allowed a maximum of two variable modifications per peptide and carbamidomethylation of cysteine residue was set as a static modification. We used a fasta file containing only the neoantigen sequences along with BFP reporter sequences and iRT sequences (Biognosys) to build the spectral library containing the heavy peptide sequences. Ion mobility calibration was performed with default selection of *Automatic selection of a run as reference IM* and only b and y ions were selected for library generation. Following heavy spectral library generation in Fragpipe, a custom R script was used to add light labeled transitions to the library for every neoantigen peptide sequence. Then, interference correction was performed by removing identical fragment ion pairs from a combined library of heavy and light neoantigens. Finally, for the library used for searching (A1101-combo), all peptides present in the synthetic peptide library were added to the DDA library and duplicates were removed. The data were analyzed in DIA-NN version 1.8.2 beta 11 at 5% precursor FDR by importing the spectral library created with Fragpipe v20. Quantification strategy was set as Robust LC (high precision), cross-run normalization as RT-dependent, and library generation as smart profiling. All of the neoantigen identifications by DIANN were cross-validated by analyzing them with Skyline (v23.1.0.268) whenever synthetic heavy peptides were available. 100 fmol of each of the previously predicted neoantigens for A*11:01 allele (total 57) were spiked into each dilution sample before data collection. The ratio of endogenous light peptide MS1 Area signal to spiked in synthetic heavy was then used to back calculate attomole amount for each neoantigen peptide.

**Table 1:** Overview of spectral library characteristics.

| **Spectral library name** | **Cell line** | **Generation** | **Number of precursors** | **Peptide length** |
| --- | --- | --- | --- | --- |
| Empirical library | A375 | empirical | 20,178 | 7-15 |
| Predicted library | A375 | predicted | 382,537 | 8-13 |
| A1101-combo | C1R -<br>A*11:01 | Empirical, plus<br>neoantigen<br>sequences | 32,830 | 7-15 |

### Statistical analysis

All data analysis was performed using custom scripts in python (3.9.4) with packages pandas (1.1.5), numpy (1.22.2), plotly (5.4.0) and scipy (1.7.3) or R (4.3.1) with packages dplyr (1.1.3), data.table (1.14.8), stringr (1.5.1), and ggplot2 (3.4.4) Statistical analysis parameters, if applicable, are described in the respective paragraph and/or figure legend.

### Experimental Design and Statistical Rationale

Experiments were performed using A375 cells or engineered C1R cell lines (see respective sections). A375 DIA experiments were performed using three technical replicates. We used the same batch of cultured cells for library creation and DIA injections. Statistical analysis parameters, if applicable, are described in the respective paragraph and/or figure legend. For dilution samples, injections were run from low input to high input to reduce carry over effects.

## Results

### HLA peptide analysis with diaPASEF workflow

The analytical depth and sensitivity of DIA make it attractive for low-yield samples, such as HLA-peptides. For the initial assessment of a diaPASEF immunopeptidomics workflow, we applied our automated immunoaffinity purification protocol to enrich HLA-I peptides from A375 multiallelic melanoma cells (Figure 1a). In short, we lysed 125 e^6^ A375 cells and enriched HLA-I peptides. Peptides from an equivalent of 75 e^6^ cells were further split into 3 replicates, fractionated, and analyzed in DDA mode. To create an experimental DDA-based spectral library, data was analyzed using Fragpipe resulting in a final library with 20,178 sequences. The remaining amount was injected in technical quadruplicates (12.5 e^6^ cells each) in diaPASEF mode (Figure 1b, Suppl. Figure 1a,b) and analyzed by DIA-NN. This led to identification of 10,297 HLA-I peptides from the equivalent of 12.5 e^6^ cells in just 60 min of LC gradient time (Figure 1c, d). Notably, we observed a high degree of data completeness with >9,500 peptides present in all 4 replicates (Supp. Figure 1c). Interestingly, > 35% of identified peptides are singly charged, which emphasizes the importance of isolation polygon optimization (Figure 1e). Furthermore, all replicates exhibited similar quantitative behavior with a median R^2^ of 0.96 and a median coefficient of variation of 12% (Figure 1f). Based on their length distribution and presence of A375 allele-typical anchor residues, we could further verify that identified peptides are indeed HLA-bound with > 92% of all identified peptides being 8-13-mers, > 70% are predicted to bind to a specific allele, and ∼5% could be assigned to multiple alleles in line with observations made for DDA based experiments (Figure 1 g).

**Figure 1:**
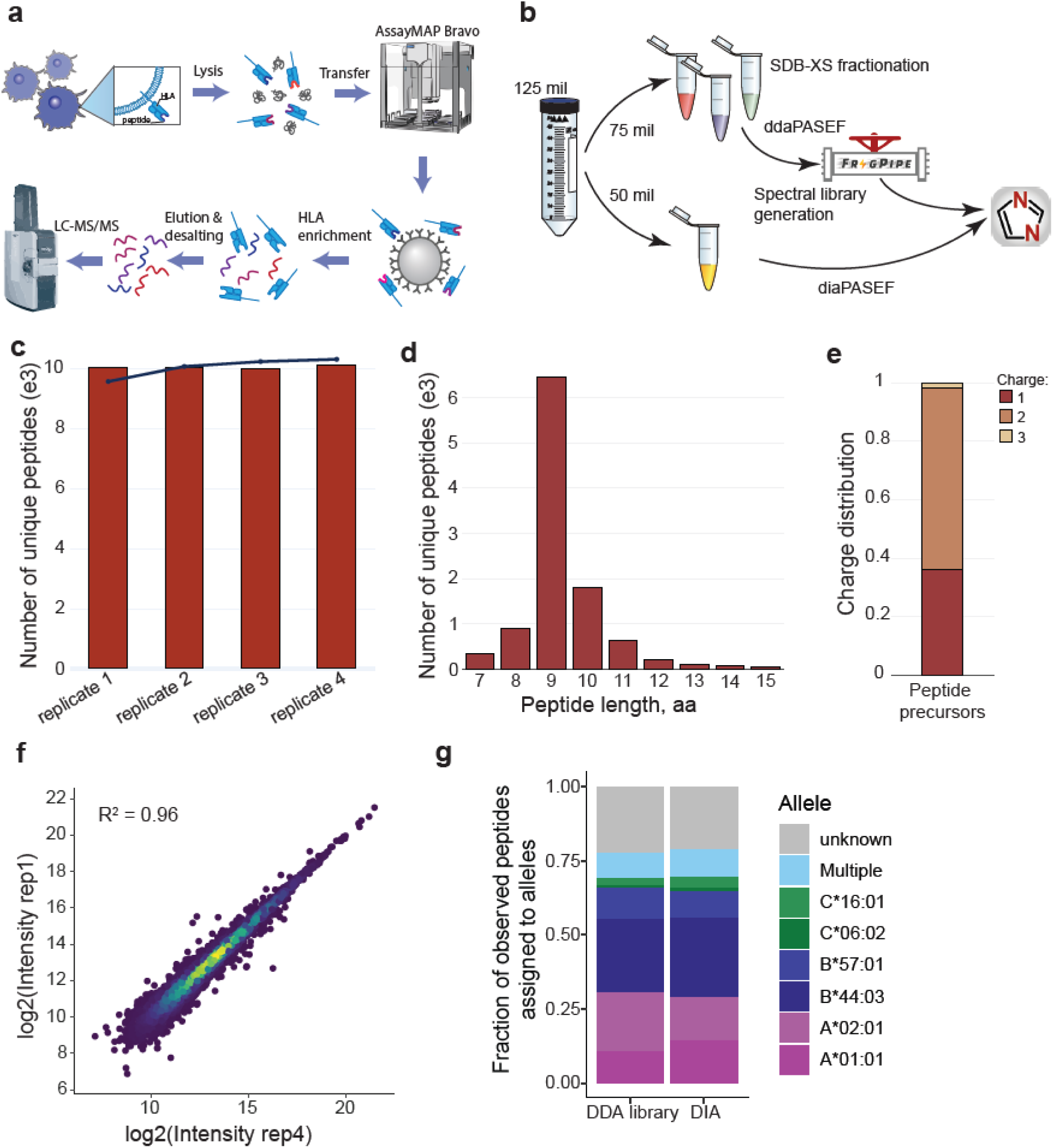
diaPASEF identifies HLA peptides with high reproducibility. a) Overview of HLA peptide enrichment workflow b) A sample specific spectral library was created from HLA peptides eluted from 3x 25 e6 cells, fractionated and acquired in ddaPASEF mode. Four replicates were analyzed in diaPASEF mode and spectra were interpreted using DIANN with the sample matched spectral library. c) Number of unique peptide identifications. d) Length distribution of peptide IDs. e) Charge distribution f) Intensity correlation between 2 replicates. g) Fraction of observed peptides assigned to alleles in DDA library and DIA peptide identifications.

### DIA offers high immunopeptidome coverage in half the analysis time

We then enriched HLA peptides from reduced cell numbers (500,000 - 10e^6^ A375 cells) in triplicate and found that DIA also increased immunopeptidome coverage when starting with fewer cells (Figure 2a). Notably, DIA identified more precursors than a previously acquired DDA dataset for all cell inputs analyzed using only a 60 min gradient compared to a 120 min gradient for our standard DDA method. Median precursor quantity increased when isolating peptides from a larger number of cells, reflecting the expected abundance changes of HLA ligands (Figure 2b). Furthermore, DIA allowed the identification of >3,000 precursors in all conditions (Figure 2c), while only >1,000 peptides were found in all samples with DDA (Figure 2d), likely due to stochastic intensity-based selection of precursors for MS/MS. For both methods, the majority of peptides are shared between the two higher input conditions (7,388 for DIA, 4702 for DDA).

**Figure 2:**
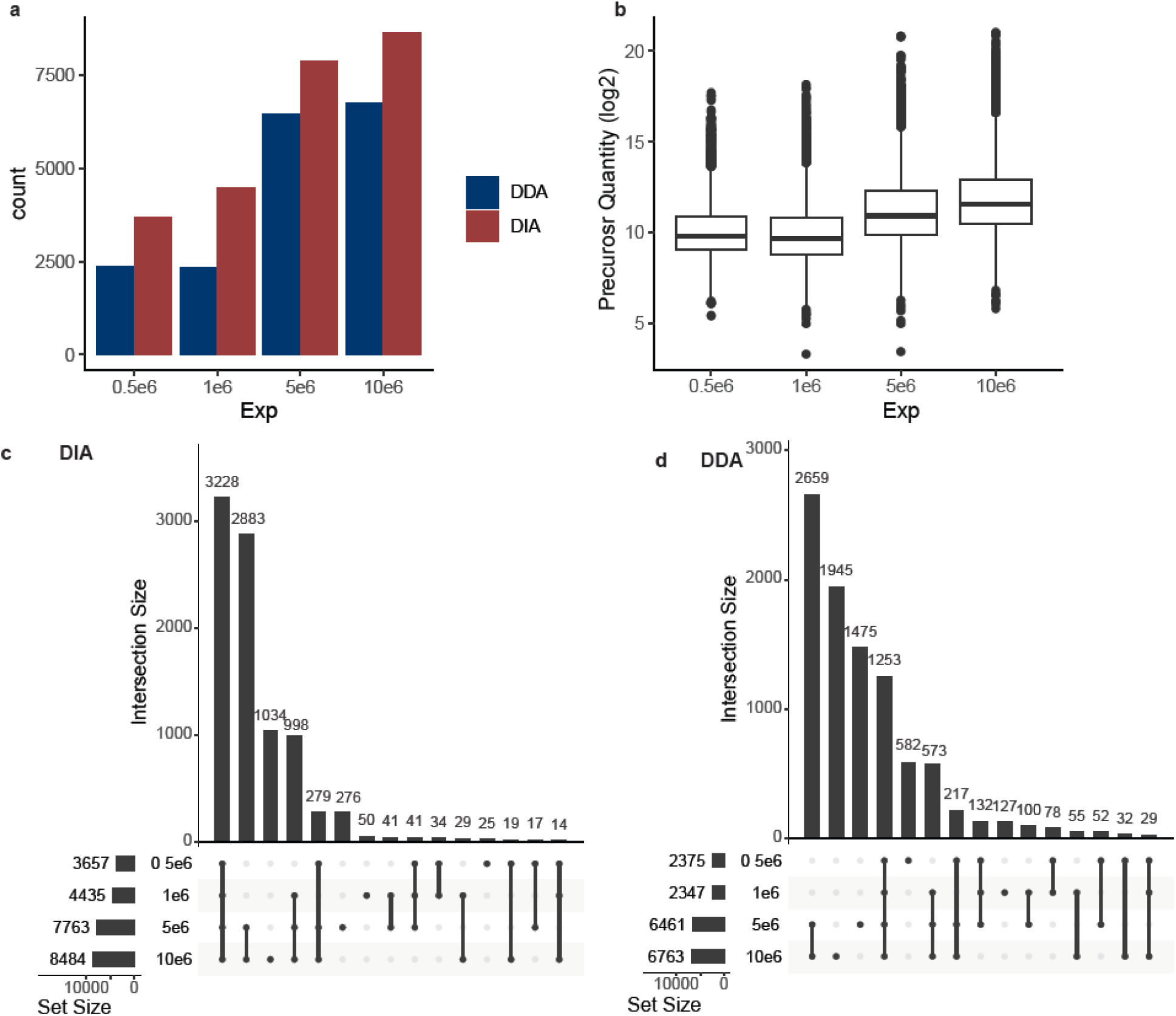
DIA outperforms DDA in half the analysis time. a) HLA peptide identifications in DDA and DIA measurements of peptides enriched from decreasing number of cells. b) Precursor quantity in DIA across samples. c) Upset plot for DIA identifications. Horizontal bars at bottom left indicate the number of total unique peptides per acquisition; dots and lines under the vertical bars identify peptides identified in only one (single dot) or multiple acquisitions (dots with lines). d) same as c) for DDA identifications.

### Predicted spectral features are a good estimate of empirically determined parameters

Experiment-specific spectral libraries facilitate a sensitive matching of HLA peptides, acquired with DIA. However, creation of such a library comes at the cost of vast material expenses and typically up to 80% of all available starting material is used to generate a deep experiment-specific spectral library. Alas, often sample specific libraries have a limited coverage due to low quantity of eluted HLA peptides and inherent disadvantages of the DDA approach. Deep learning algorithms for prediction of MS/MS spectra for existing peptide sequences offer an alternative approach for creation of experiment-specific sample libraries. However, due to the non-tryptic nature of HLA peptides, the number of theoretical 8-13-mers from the human proteome exceeds 50 million variants, which complicates directDIA searches.

We, therefore, compared a library containing predicted spectral features of sequences in the library and the empirically generated spectral library. First, to verify the accuracy of retention time prediction, we inspected the correlation between predicted retention time (RT), calculated by DIA-NN, and measured RT (Supp. Figure 2a, b). Our predicted library showed a slightly higher deviation in comparison with the empirical library that might point to bigger differences in mean RT prediction between libraries. Indeed, we observed that standard deviation for difference between measured RT and predicted RT in predicted library is ∼3-fold higher than in empirical (Figure 3a). However, the difference of DIA-measured RT apex of identified elution profiles between two libraries was within 3 seconds meaning that both libraries managed to identify the same elution profiles (Figure 3b). We next asked if the predicted spectral library contains sufficient information for quantifying fragment ions in DIA files. Reassuringly, the predicted library yielded a similar fraction of quantified fragment ions hinting on a good quality of fragment intensity predictions by Salud (Figure 3c). Moreover, the overall quality of predictions (median spectral angle = 0.68) was on par with the experiment-specific library (spectral angle = 0.7) (Figure 3d). In concordance with previous studies (6), we found a drop of ∼15% in the number of identified HLA peptides with a predicted library in comparison with empirical library (8,424 vs 10,597). More than 85% of peptides, identified with predicted spectral library, were shared with the empirical one. Interestingly, we found ∼15% (1,062 peptides) to be found only with predicted library while 38% of peptides were unique to the empirical library (Figure 3e).

**Figure 3:**
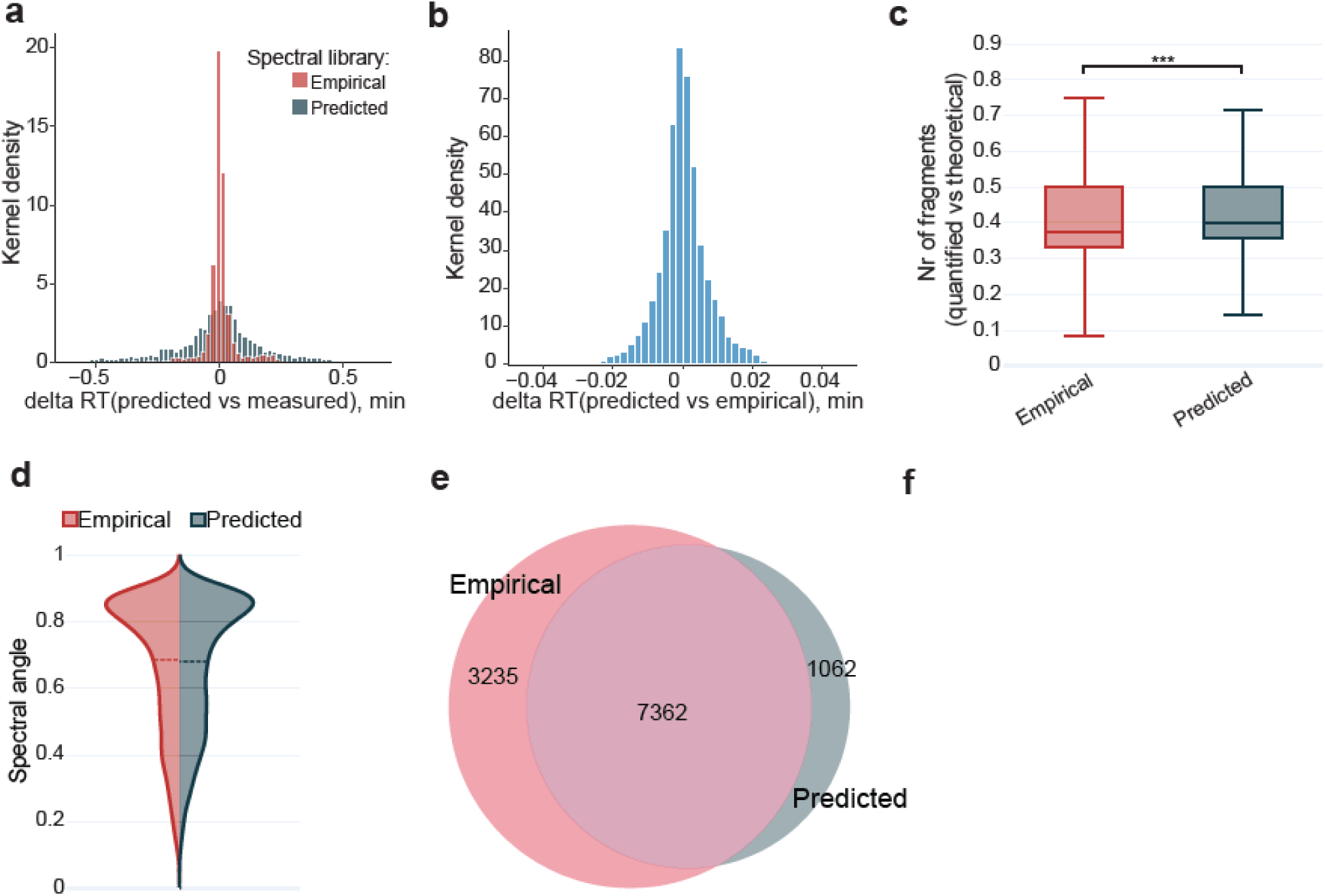
Predicted spectral library for HLA peptides can be used to query diaPASEF data. a) Comparison of empirically observed retention times to predicted retention times of peptides in the spectral library. b) Difference of DIA-measured RT apex between both libraries. c) Ratio of quantified fragment ions compared to available fragment ions in empirical and predicted libraries. *** denotes p value <0.01. d) Spectral angle of identifications in empirical and predicted libraries. Dashed line indicates the median spectral angle. e) Overlap of identifications when using empirical or predicted libraries.

### High-sensitivity immunopeptidomics with predicted libraries

Encouraged by the correlation of empirical and predicted spectral library we generated a twofold prediction strategy to create a synthetic spectral library for A375 that comprised all possible 8-13-mer binders. First, human canonical proteins were predicted for presentation (343,034,958 peptide-HLA allele combinations) using the state of the art predictor HLApollo(13), reducing the possible search space to only 382,537 sequences that could possibly be presented. Retention time, fragment ions, and ion mobility were then predicted for this reduced set of possible precursors using Salud to generate an allele-specific, sample matched spectral library (Figure 4a). The predicted library was able to identify similar or more peptides for samples enriched from 5e^5^ - 25e^6^ cells summarized across triplicates (Figure 4b). Both approaches resulted in high degree of reproducibility with the median coefficient of variation < 0.2 across all conditions (Figure 4b). For example, at the 25e6 input level we identified > 12,643 peptides with in-silico library versus ∼8,288 peptides with the DDA-based library. We observed ∼ 70% overlap of peptide IDs between two libraries at 25e^6^ input while at 5e^5^ input it decreased to ∼60% (Figure 4c). Interestingly, at an input amount of <5e6 cells, we identified more peptides with the empirical library than in the *in silico* library analysis. However, when comparing motifs of overlapping and unique peptides to either library for the 0.5e6 experiment, we found that all peptides show characteristic sequences for A375 presented peptides (Figure 4d). Upon examination of additional quality criteria such as intensity and spectral angle, peptides identified by the empirical library but not the predicted library show lower intensity and lower spectral angles and might be less confident identifications to begin with (Figure 4e, Supp. Figure 4 a-b).

**Figure 4:**
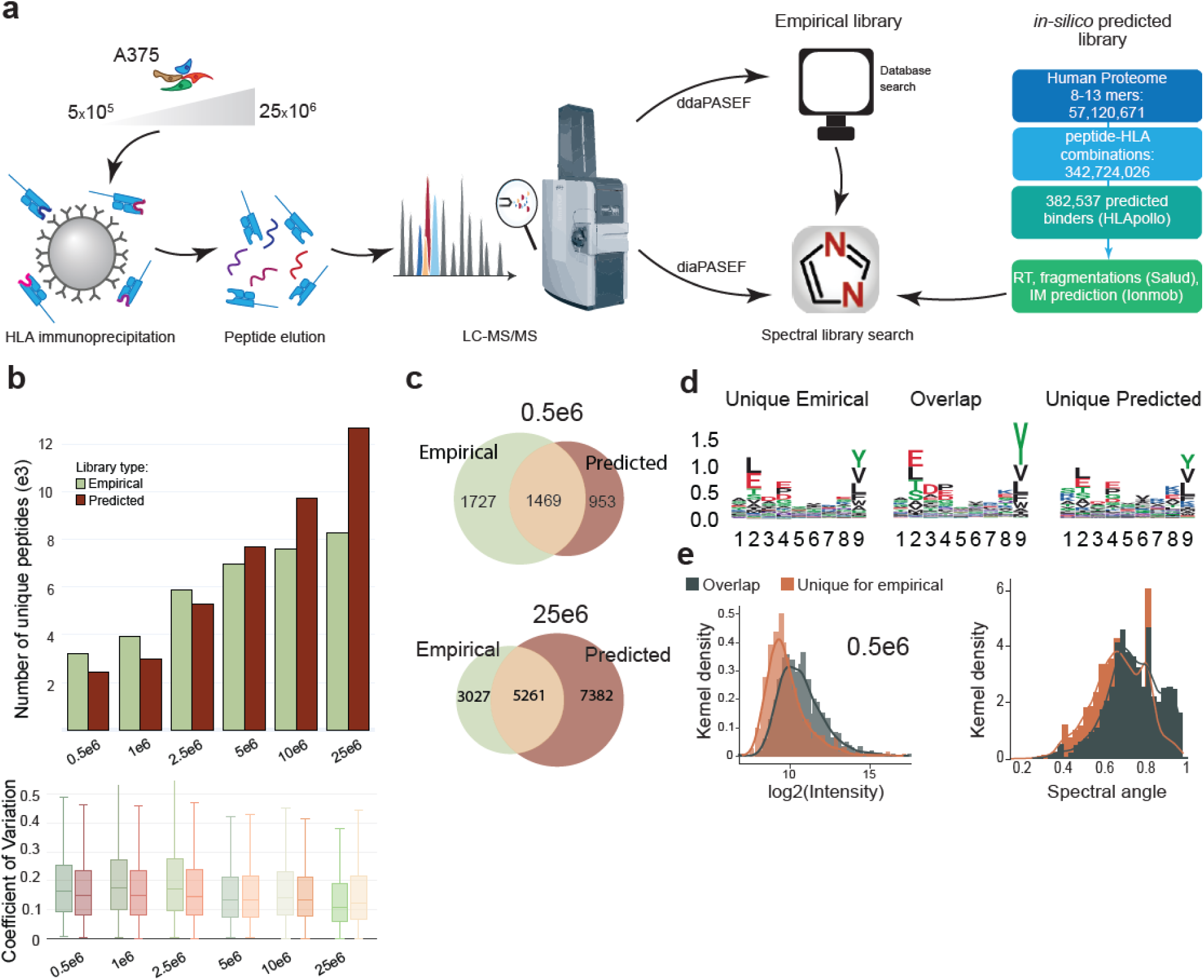
Predicted spectral library for all A375 HLA alleles generally outperforms empirical library. a) In silico library creation by predicting all possible HLA-peptide combinations for A375 cells and spectral feature prediction with Salud. b) Number of unique HLA peptides identified with empirical (green) or predicted library (red). c) Venn diagrams of identified peptides for 1e6 (top) and 25e6 cells (bottom). d) Coefficient of variation of identifications using different spectral libraries. e) Intensity correlation between empirical and in silico matches. f) Low abundant precursors are preferentially identified with the empirical library only. g) Spectral angle for identifications overlapping between libraries or uniquely identified with empirical library.

### Ion mobility reduces co eluting peptides

Using the predicted spectral features of over 300,000 possibly presented peptides, we further evaluated the ion distribution across DIA windows and LC gradient (Supp. Figure 4a). The majority of predicted precursors fell into windows 1.2 (1/k0:0.4-0.87; m/z: 300.53 mz-430.23 mz) and window 12.1 (1/k0: 1.09 - 1.7; m/z: 975.48-1340.62) with >200 precursors eluting in the middle of the LC gradient (Supp. Figure 4b). However, these are deliberately large windows and not many actual features are observed for these mass / ion mobility combinations. All other DIA window and PASEF cycle combinations contain fewer than 50 precursors that are predicted to be analyzed at a given time in the same window. To characterize the extent of co-fragmentation on HLA peptide sequences, we calculated the occurrence of shared peptide fragments in a PASEF cycle window at a given predicted LC elution point. We compared fragment ions in a wide m/z window and found that many of the predicted, coeluting ions share fragment ion masses wouldn’t be uniquely assigned to a particular peptide unless the ions are of longer length (b/y^9^ or higher, Supp. Figure 4c). Reassuringly, this effect is drastically reduced for windows with fewer co-eluting features (such as 8.2), where only lower m/z fragment ions (<b3/y3) would be shared among coeluting precursors and the majority of fragment ions can be uniquely assigned to a single precursor (Supp. Figure 4c). Overall this analysis shows that the addition of ion mobility significantly reduces the problem of indistinguishable, coeluting HLA peptides in DIA.

### DIA quantification benchmarking with SILAC

Next, we derived a SILAC HLA peptide DIA strategy to evaluate quantitative accuracy and precision. A375 cells grown in heavy SILAC (K, R, L) and light medium were mixed to a total of 20e^6^ cells each at ratios 1:1, 1:2, 1:4, 1:9 and 1:19 prior to lysis, HLA peptide enrichment and DIA (Figure 5a). DIA data was analyzed using an A375 spectral library and DIANN with SILAC specific parameters (Methods). Not all presented peptides contained an amino acid used in the SILAC mix. Therefore, H/L ratios could be calculated for 70-80 % of peptides identified in each experiment, with 1,570 peptides quantified across all 5 ratios. Median H/L ratios for all peptides identified per sample closely matched expected H/L ratios for 1:1, 1:2 and 1:4 dilutions with only 0-4% relative error (Figure 5b). For dilutions 1:9 and 1:19, measured ratios had 16% or 46% relative error compared to the expected ratio. This is likely due to identification of lower intensity precursors, mis-identifying light labeled peptides as heavy labeled and vice versa (Supp. Figure 5a). Indeed, when examining the quality of the spectral match, heavy identifications have a greater C-Score distribution than light labeled peptide identifications, with a median of 0.5 C-score for heavy peptides in the 1:19 dilution (Supp. Figure 5b). However, while additional filtering for channel and translated q values(16) improved C-scores of remaining peptides (Supp. Figure 5c), it did not affect median ratio or remove peptides with extreme ratios (Supp. Figure 5d). This indicates that peptides are either misidentified or suffer from high signal to noise at extreme ratios and need to be evaluated carefully. Overall, quantitation is accurate up to 5-fold dilution, in line with similar studies on low input samples(17).

**Figure 5:**
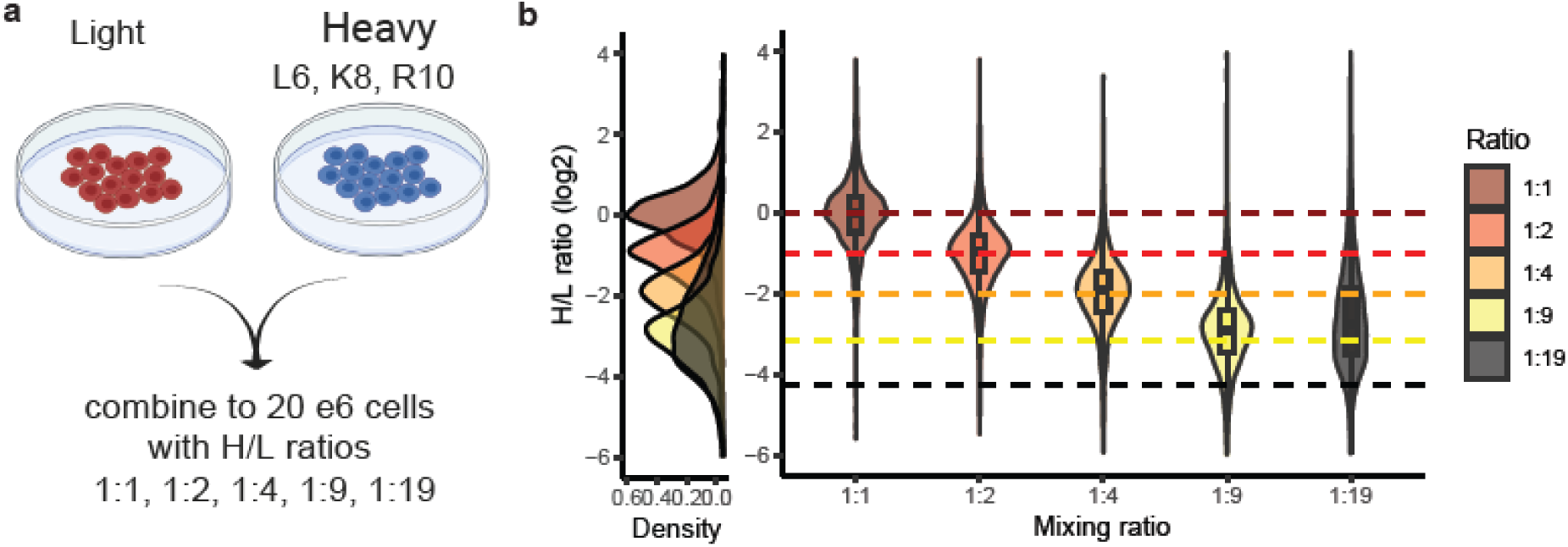
diaPASEF quantification benchmarking with SILAC. a) SILAC labeling strategy for MHC peptide analysis. b) Violin and density plot for log2 H/L ratios across mixing ratio experiments. Box plots show median ratio, dotted lines indicate expected ratio.

### Identification and quantification of common cancer neoantigens

Next, we evaluated how DIA analysis could benefit neoantigen identification and quantification. We used a mono-allelic C1R cell line expressing HLA-A*11:01 that was transfected with a neoantigen cassette encoding 47 common cancer mutations (Figure 6a)(15). HLA complexes were enriched from 1e^6^ - 50e^6^ cells and eluted, followed by DDA for library generation and then DIA. Prior to DIA analysis, 100 fmol of synthetic, heavy standard peptides were spiked into the sample. This allowed for an internal control and helped eliminate false positive identification as spiked in peptides co-elute and co-fragment. We observed a steady increase in identifications up to 10e6 and 25e6 cells. Interestingly, identifications decreased for 50e6 cells in DIA runs compared to DDA, likely due to space charging, gradient or library constraints (Figure 6b). We observed an increase in abundance of neoantigens while the spike-in MS1 area stayed relatively constant (Figure 6c). For example, while a neoantigen from EGFR G719A was low abundant at 1e6 cell input, it could be detected across all input ranges tested with increasing presentation on the surface (Figure 6d, e). We identified 16 peptide sequences from the neoantigen construct, the majority of sequences from KRAS mutant sequences. As expected, DIA provided a more complete coverage across the dilution series and detected 14/16 neoantigens in as low as 1e6 cell input (Figure 6f). Using the signal of heavy spike-in peptides, we quantified the relative abundance of 13 neoantigens on the surface starting at 33 attomoles for 1e6 cells. For comparison, 20 neoantigens were found in a targeted PRM experiment and 9 with DDA from 166 e6 cells(15). This highlights the general utility of DIA acquisition for quantification in immunopeptidomics experiments when following selected peptides of interest across samples.

**Figure 6:**
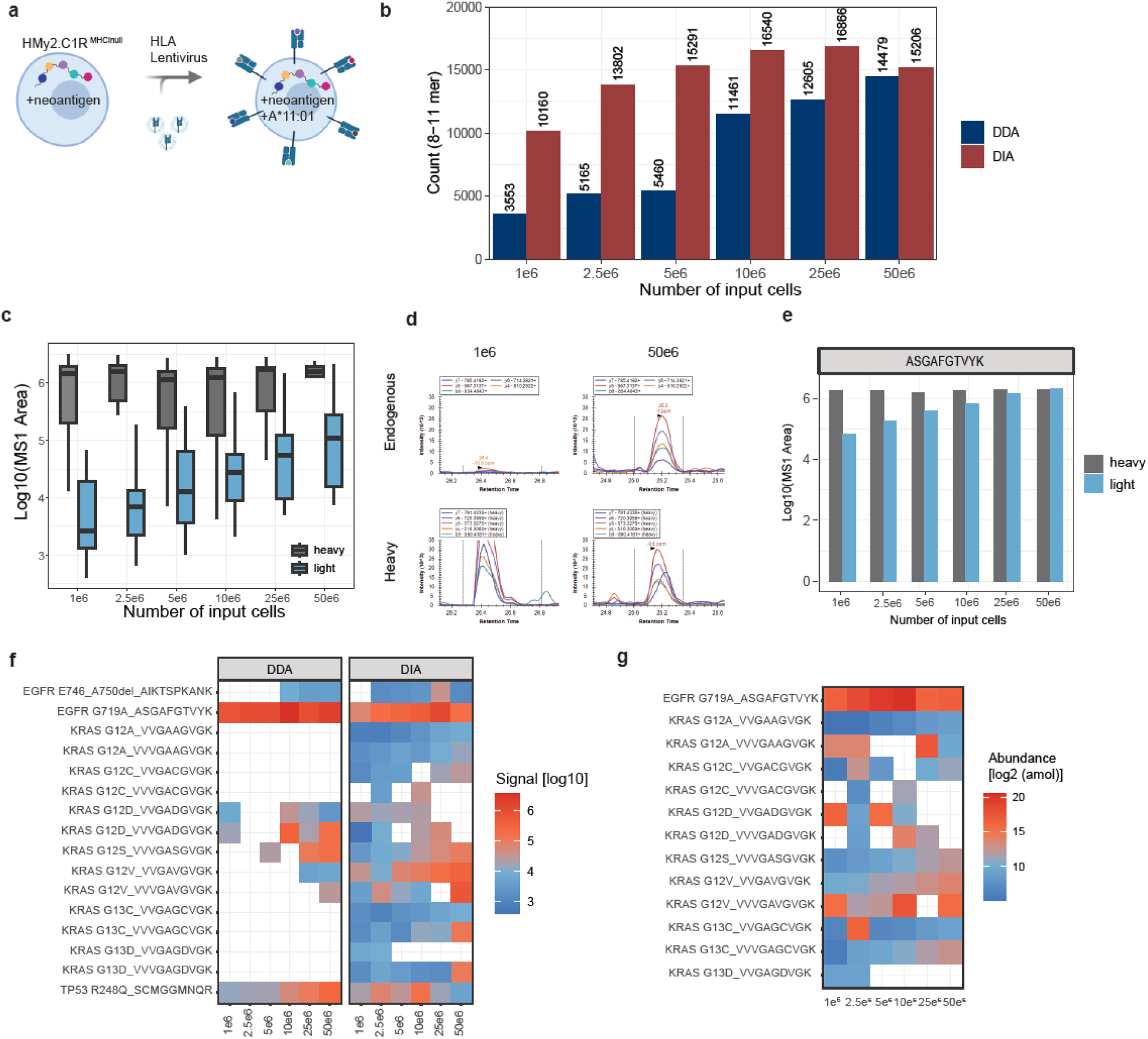
diaPASEF identification and quantification of A*11:01 neoantigens. a) MHC-I null C1R cell lines were transfected with a neoantigen cassette containing 47 common cancer mutations and a MHC-I allele of interest and MHC ligands were eluted from increasing cell amounts. b) MS1 area of endogenous presented peptides increased while synthetic spike in areas were stable. c) Extracted Ion Chromatograms (XIC) of endogenous (top) and heavy labeled neoantigen peptide. d) Same as c but quantified using DIANN MS1 Area. e) Intensity of neoantigen identifications in DDA and DIA acquisition increases with increasing sample input. f) Abundance of all neoantigens identified and quantified in A*11:01 C1R cells normalized to 100fmol of heavy synthetic spike in peptides.

Collectively, these results point to high analytical depth and exceptional quantitative reproducibility of DIA in the context of immunopeptidomics.

## Discussion

Advancements in instrumentation have further increased the application of DIA in numerous proteomic studies. Key advantages include reproducibility between replicates and relative quantification across a large number of samples. For DDA based immunopeptidomic measurements, reproducibility between replicates historically ranges between 60-70%. With DIA we achieved over 99% overlap of identified peptides in replicates. The integration of ion mobility, such as FAIMS or TIMS, decreases the number of co eluting precursors and fewer overlapping or shared fragment ions allow for higher confidence in identifications.

Though DIA resulted in great depth of coverage with reduced acquisition time of an immunopeptidome sample of interest, additional sample material is needed to create a spectral library. Due to the biological complexity of HLA peptidome samples, library based approaches are necessary to keep FDR low. Other studies have predicted a smaller set of peptides of interest to be included in a spectral library, generate a large library on available public HLA peptide data on either sequence or spectral level independent of sample context and HLA types present in a sample(6, 7). We propose a two-step strategy that first employs state-of-the-art HLA binding predictors to reduce the possible number of peptide sequences in the human proteome and tailor them to the sample of investigation. The second step includes the prediction of spectral features similar to published approaches. This combined prediction approach leads to similar or more identifications depending on the amount of peptide loaded on column. It can be further refined by adding contaminant species and non-canonical peptide sequences. Prediction results can also be used to filter peptides further, for example, peptides uniquely identified in experimentally determined spectral libraries are of lower intensity and show poorer spectral angle, indicating that these might be lower confidence hits.

Using all possible presented precursors, we could also evaluate our data acquisition approach. We found that ion mobility in general reduces the number of co eluting and co fragmenting peptides and even for highly similar peptide sequences like HLA presented peptides, most DIA spectra contain <50 features with distinct fragment ions. Additional data acquisition approaches further improving window design, as well as novel approaches such as sliceDIA or synchroPASEF(18, 19), midiaPASEF(20), and narrow window acquisition(21) can further decrease co-elution and co-fragmentation leading to confident identifications.

DIA acquisition is quite powerful for quantification across many samples compared to DDA based label free or TMT-based quantification. For instance, assessing peptide abundance changes due to varying treatment conditions or input amounts in a consistent sample background is likely executed more efficiently with DIA. But due to the inherent diversity of the HLA peptidome, stratification by respective HLA types is pivotal before conducting such experiments. We evaluated quantitative accuracy using a SILAC approach and found that DIA analysis can accurately recover known quantities up to a 5 fold difference. Quantification of larger ratios suffers from background noise contamination or incomplete fragmentation and wrong assignment of spectral matches. This is in line with reports in other studies including single cell analyses(17, 22). In cell line samples carrying common tumor mutations, we were able to detect neoantigen peptides as low as 1e6 cells input into the IP and quantify increasing amounts of peptides with increasing amounts of cells. If coupled with a hip-MHC approach(23), ratios of light to heavy peptides can be used to determine the copy number of peptides presented on the surface, while still gathering data on all other peptides present in the sample, and hence not losing additional peptidome information discarded in PRM acquisitions(24, 25). In summary, the integration of DIA with ion mobility emerges as a potent data acquisition approach. It enhances reproducibility and facilitates effortless quantification of peptides within the same HLA allele type background. Additionally, the library-free strategy we proposed for HLA immunopeptidome measurements can be conveniently constructed using existing resources. This approach, coupled with DIA measurements, ensures robust quantification which can significantly increase throughput, proving advantageous to studies appraising immunotherapy mode of action.

## Acknowledgements

The authors want to thank Felipe da Veiga Leprevost, Meena Choi and Alessandro Ori for insightful discussions.

## Author contribution

CMR and SK conceptualized the study; DO, HG, KL performed experiments; ZZ, CMR performed library prediction analyses; DO, HG, SK curated and analyzed data and wrote the manuscript with input from all authors.

---

**Supp. Figure 1.**
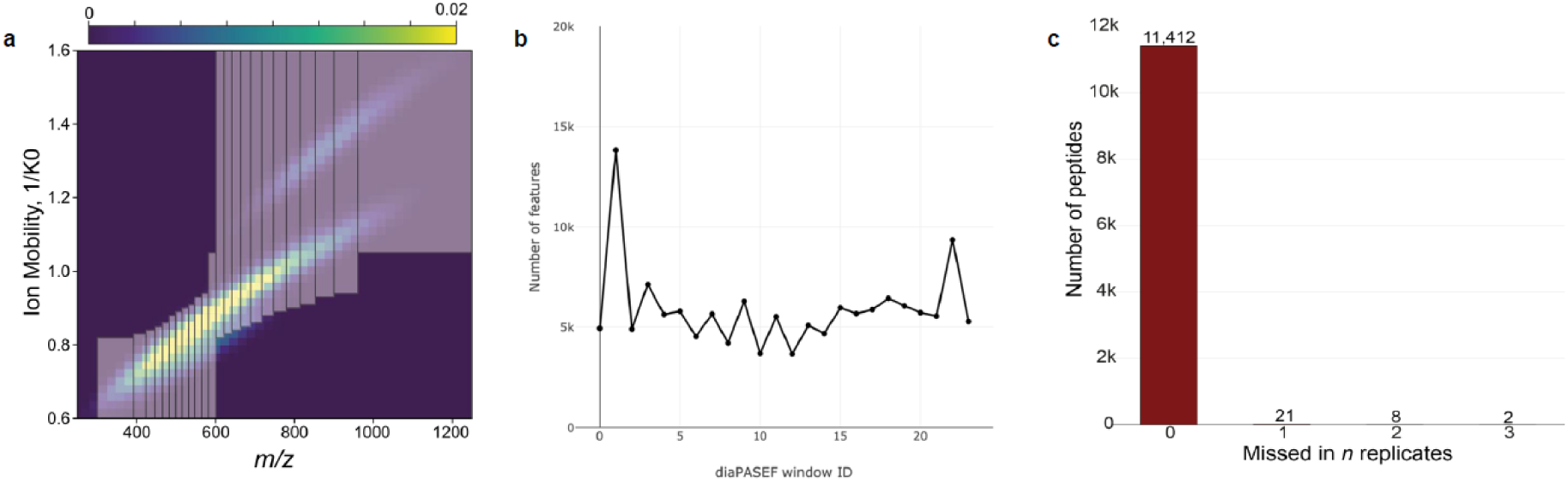
a) Variable window design for diaPASEF with MHC Class I peptides. b) Variable window design enables equal feature distribution per cycle. c) Number of peptides missing in replicate injections.

**Supp. Figure.**
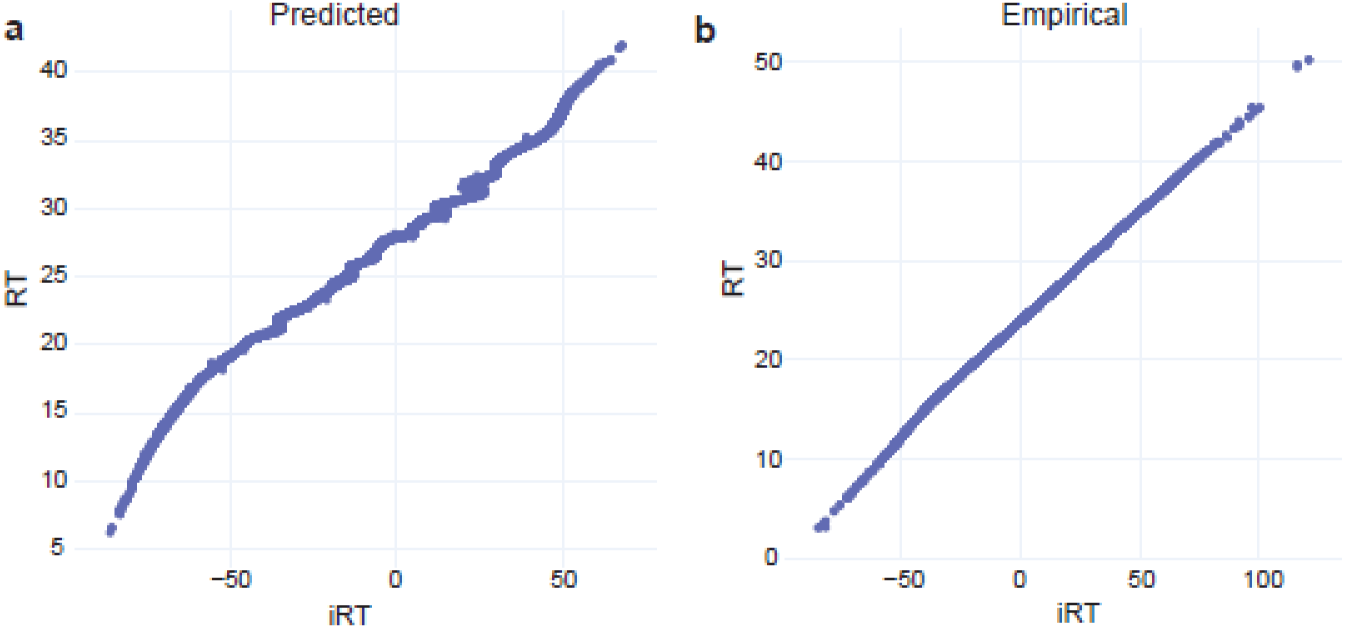
2 Correlation between predicted retention time as calculated by DIA-NN (iRT) and measured RT for Salud predicted libraries (a) and empirical library (b).

**Supp. Figure 3.**
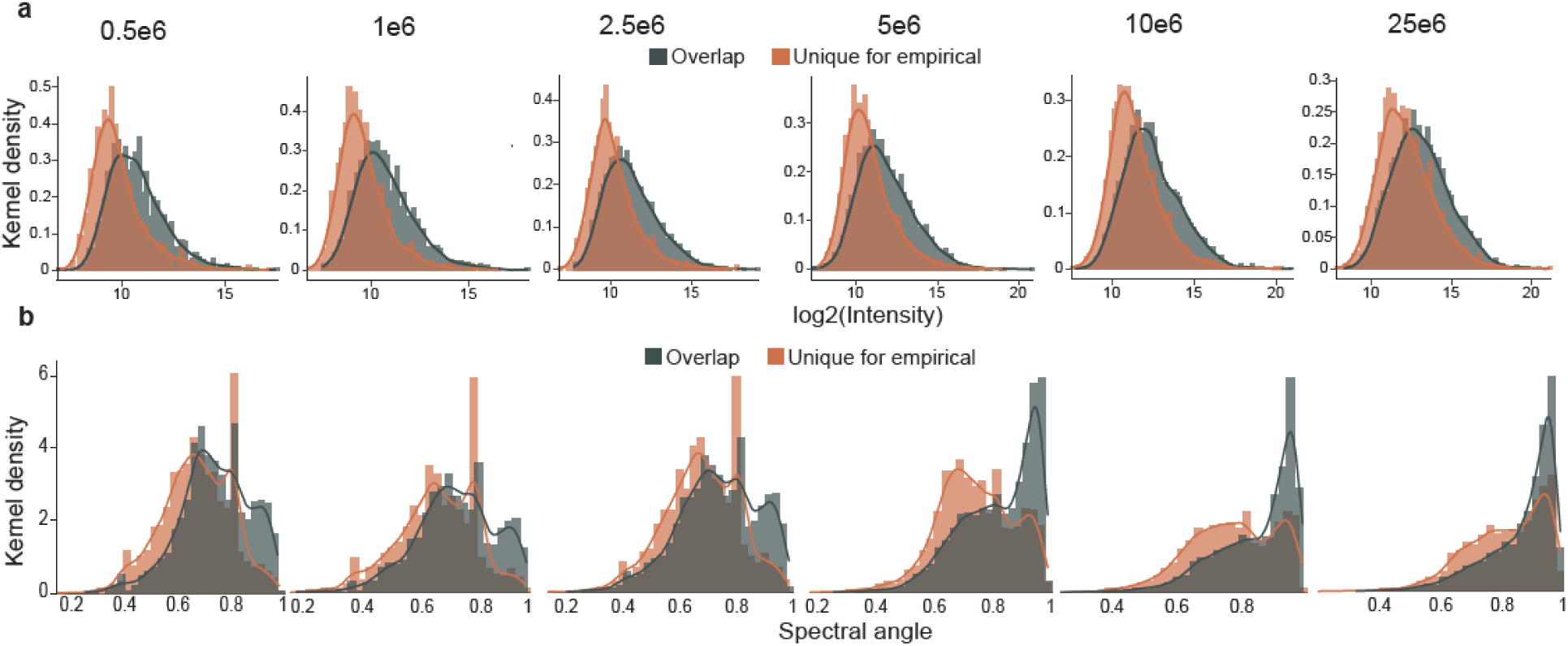
a) Low abundant precursors are preferentially identified with the empirical library only. b) Spectral angle for identifications overlapping between libraries or uniquely identified with empirical library.

**Supp. Figure 4.**
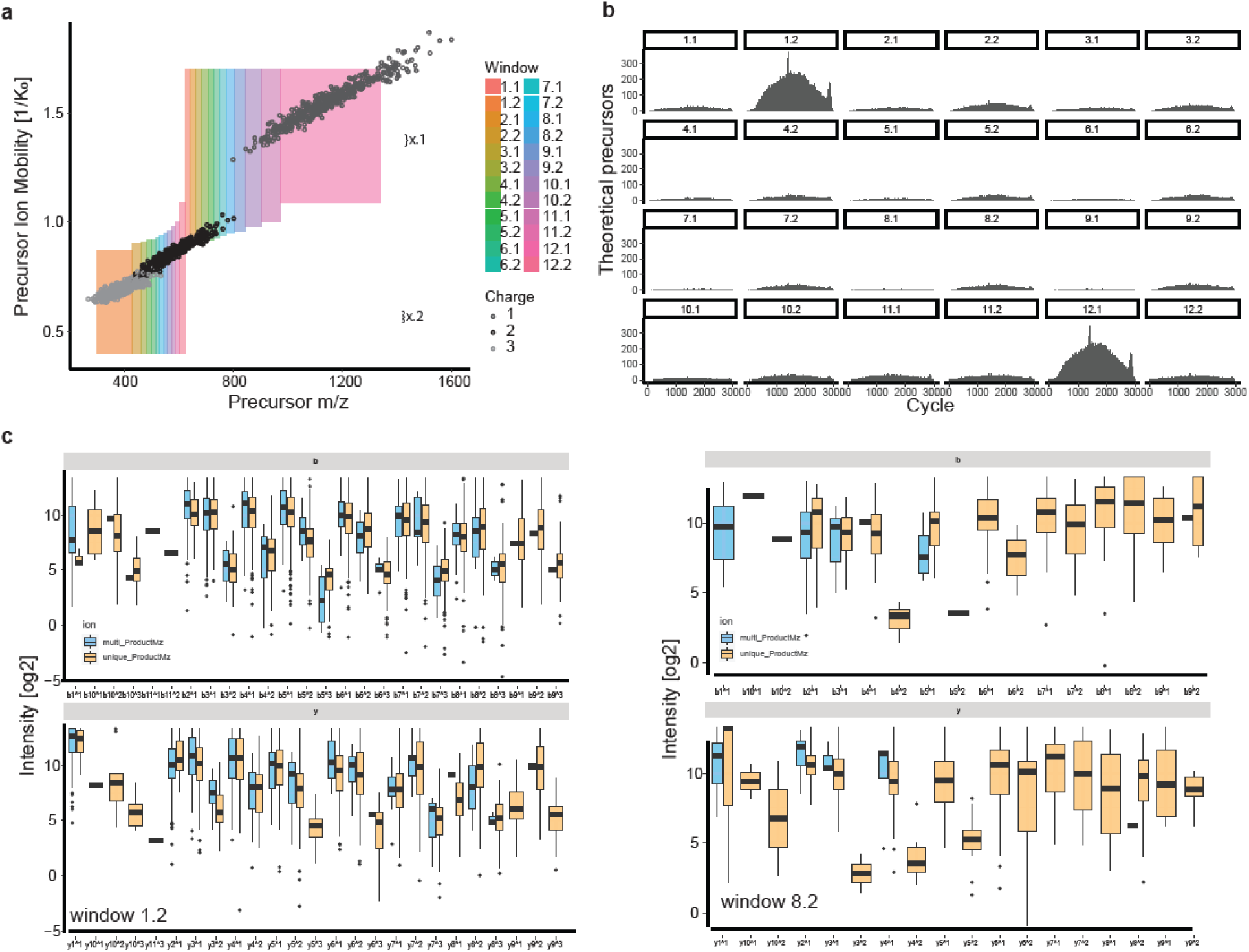
a) Predicted Ion Mobility for theoretically presented peptides, squares indicate elution window during acquisition. b) Number of theoretical eluting precursors per ion mobility window and instrument acquisition cycle. c) Fragment ion distribution of all possible coeluting predicted precursors in large window of mostly singly charged precursors (1.2) and a narrow window of doubly charged precursors (8.2). Color indicates fragment ion occurences in multiple precursors (blue) or in a unique precursor (orange) in a particular window/cycle combination.

**Supp. Figure 5.**
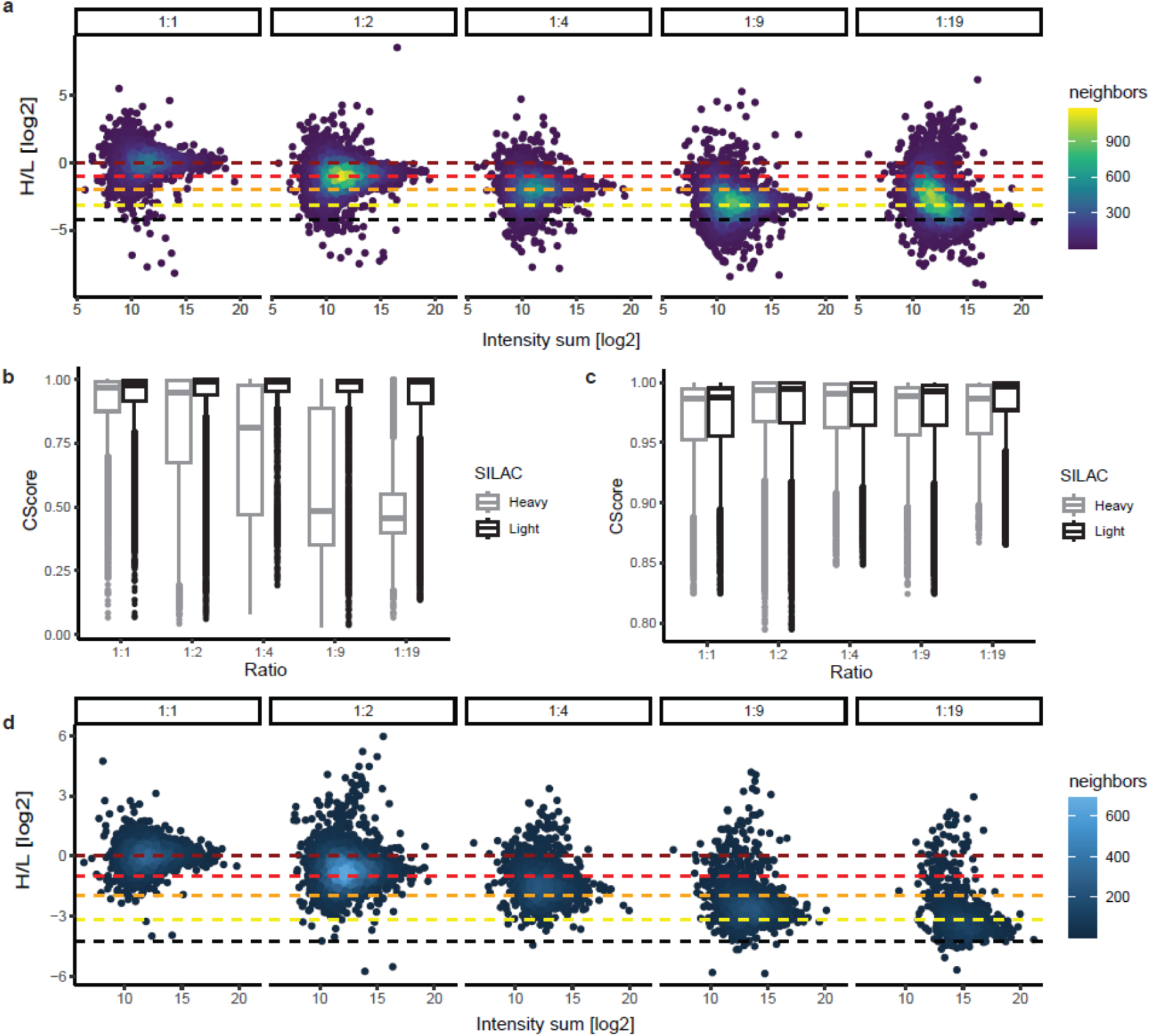
a) 2D density plots showing log2 H/L ratios for peptides according to their summed H+L intensity. b) C-Score distribution of heavy and light peptide identifications for each mixing ratio. c) Same as b, but after filtering for channel q-vale <0.01 and translated q-value <0.01. d) H/L density based on summed intensity across mixing ratios after filtering for channel q-vale <0.01 and translated q-value <0.01. Dotted lines indicate expected ratio.

